# Aggregation promoting sequences rather than phosphorylation are essential for Tau-mediated toxicity in *Drosophila*

**DOI:** 10.1101/2024.12.22.629946

**Authors:** Amber Cooper, Bradley Richardson, Eva Ruiz Ortega, Yongrui Zhang, Ben Batchelor, Aarya Vaikakkara Chithran, Jie Liu, Tianshun Lian, Miguel Ramírez Moreno, Benjamin Boehme, Leila Abtahi, George Devitt, Lovesha Sivanantharajah, Efthimios M. C. Skoulakis, Douglas W. Allan, Amritpal Mudher

## Abstract

**Background:** Disease-modifying therapies for tauopathies like Alzheimer’s disease have targeted Tau hyperphosphorylation and aggregation, as both pathological manifestations are implicated in Tau-mediated toxicity. However, the relative contributions of these pathology-linked changes to Tau neurotoxicity remain unclear.

**Methods:** Leveraging the genetic tractability of *Drosophila*, we generated multiple inducible human Tau transgenes with altered phosphorylation status and/or aggregation propensity. Their individual and combined impact was tested *in vivo* by quantifying Tau accumulation and neurodegenerative phenotypes in the aging fly nervous system.

**Results:** We report that phospho-mimicking Tau (hTau2N4R^E14^) induced profound neurodegeneration, supporting a neurotoxic role for phosphorylation. However, when we rendered hTau2N4R^E14^ aggregation incompetent, by deleting the ^306^VQIVYK^311^ motif in the microtubule-binding region, neurotoxicity was abolished. Moreover, a peptide inhibitor targeting this motif efficaciously reduced Tau toxicity in aging *Drosophila*.

**Conclusion:** Neurodegeneration mediated by Tau hyperphosphorylation is gated via at least one aggregation-mediating motif on the protein. This highlights the primacy of blocking Tau aggregation in therapy, perhaps without the need to clear phosphorylated species.

## Introduction

The presence of highly phosphorylated, insoluble inclusions of the microtubule associated protein Tau (MAPT) within neurons and/or glia of the central nervous system (CNS) is a common pathological hallmark of Alzheimer’s disease (AD) and other neurodegenerative dementias collectively known as tauopathies(1–3). Tau hyperphosphorylation and aggregation are the two most prominent pathogenic hallmarks that influence Tau-mediated toxicity and have therefore become the primary targets for modern Tau-based diagnostic biomarkers and therapeutics(4, 5). However, the extent of their interdependence or relative impacts in the progression of Tau-mediated toxicity, neuronal degeneration and disease progression remains unclear.

Physiologically, Tau phosphorylation is implicated in the regulation of Tau-microtubule binding(6), promoting microtubule assembly, stabilisation and spacing of the microtubule network within neurons(7). However, a hallmark of the pathogenic process is Tau becoming increasingly hyperphosphorylated(8), resulting in less efficient binding to microtubules and the induction of toxic consequences at the molecular, subcellular and cellular level. *In vitro* studies have demonstrated that expressing phospho-mimicking Tau in neuronal cells has a profound effect on neuronal structure and function, compromising cytoskeletal integrity(9, 10), suppressing neuronal excitability(11) and inducing significant changes to the normal subcellular localization and distribution of Tau(11–13). Similar deficits are reported *in vivo* in *Drosophila* models of Tauopathy where driving hyper-phosphorylation, by co-expression of the Tau kinase GSK-3β, or emulating hyperphosphorylation by expressing the TauE14 transgene disrupted axonal transport(14), reduced microtubule binding(15) and induced significant neuronal, and synaptic degeneration(9, 14, 16–19). In the TauE14 transgene key disease-associated serines and threonines have been substituted with glutamic acid, with the charge on the latter emulating phosphorylation. These manipulations impair neuronal function in multiple ways; disrupting axonal transport(14), reducing microtubule binding(15) and inducing significant neuronal, and synaptic degeneration(9, 14, 16–19). Moreover, expression of TauE14 resulted in greater accumulation of Tau in sarkosyl insoluble cellular fractions and the formation of *bona fide* Tau aggregates within fly brains(17, 20, 21). Accordingly, in FTD mouse models, transgenic Tau variants can misfold and form pre-tangle-like fibrils, which can be significantly reduced with GSK-3β inhibitor treatment(22–28).

Several species of aggregated Tau have been identified in both human brain and animal Tauopathy models. However, the smaller hyperphosphorylated oligomeric aggregates are thought to mediate toxicity(29, 30). The hypothesis that aggregation of these smaller Tau oligomers mediates Tau toxicity is well-supported in cultured cell models, presenting with impaired synaptic function(13), cellular damage(31), disrupted cellular trafficking including axonal transport(32), and induction of inflammatory responses(33). In addition, transgenic animals over-expressing aggregation-prone Tau mutants and isoforms incur significant neuronal degeneration paired with cognitive and locomotor deficits(15, 34–37). Such aggregation-induced toxicity has been attributed to two hexapeptide motifs, PHF6* (^275^VQIINK^280^) and PHF6 (^306^VQIVYK^311^)(37), when these hexapeptides are deleted, or blocked by site-specific peptide-based drugs, cognitive deficits and reduced lifespan are largely ameliorated in transgenic animals(38–40).

Evidence implicates Tau phosphorylation in promoting aggregation, but the precise molecular mechanism underlying this interplay is not clear. Nonetheless, phospho-epitopes of Tau influence its aggregation, either directly or by promoting phosphorylation at additional sites(17, 41–46). The consensus view is that phosphorylation at residues that reduce microtubule binding can encourage aggregation by enhancing the local cytosolic concentration of free Tau, thus encouraging interaction with aggregation-promoting cofactors(47). Alternatively, Tau phosphorylation may induce misfolding into seed-competent and aggregation prone conformations(48, 49). Additionally, phosphorylation at certain sites appears to suppress further phosphorylation and aggregation(45, 50–53). Conversely, inhibiting aggregation through ^306^VQIVYK^311^ ablation and peptide-based drugs leads to reduced phosphorylation in *in vitro* and *in vivo* models of tauopathy(38, 39, 54, 55), suggesting that aggregation may in fact promote phosphorylation. However, such incongruency may arise from utilization of different Tau isoforms in diverse experimental systems.

Beyond the potential causal and perhaps reciprocal relationship between Tau aggregation and phosphorylation, interpreting the relative contribution of each pathological modification towards Tau-mediated toxicity and neuronal dysfunction is challenging, as it is difficult in a single model to independently and combinatorically manipulate phosphorylation, and aggregation, Tau expression levels and neuronal age, all of which contribute to degenerative phenotypes(34, 56, 57). To resolve this conundrum, a systematic assessment of the aggregation-phosphorylation interplay in a well-controlled *in vivo* experimental model is needed.

To address this problem, we assessed the relative contributions of phosphorylation and aggregation on Tau-mediated degeneration using transgenic *Drosophila*. A number of novel human Tau (hTau) mutants were generated to determine toxicity of wild type (WT) and phospho-mimicking mutants (hTau2N4R^E14^), and mutants with deletion of the aggregation-promoting Δ^306^VQIVYK^311^ sequence (hTau2N4R^DEL^ and hTau2N4R^E14/DEL^), to enable assessment of the toxicity of phosphorylated Tau which cannot aggregate. We report that the expression of WT hTau2N4R induced significant intracellular Tau accumulation and neuronal degeneration, which was greatly exacerbated by the phospho-mimicking hTau2N4R^E14^. However, removal of the ^306^VQIVYK^311^ sequence completely abolished the toxicity of the phospho-mimicking Tau, demonstrating that whilst hyperphosphorylation of Tau drives Tau-toxicity, this absolutely requires the aggregation promoting Δ^306^VQIVYK^311^ domain.

## Results

### Phosphorylation-dependent somatodendritic and axonal degeneration requires aggregation

To investigate the relative contributions of hyperphosphorylation and aggregation to neuronal degeneration, we first generated a panel of 10xUAS-hTau2N4R transgenes with specific mutation(s) and a single copy was integrated into the *attP40* locus of the fly genome. Thus, the mutants were expressed at similar levels in all experiments (Supplementary Fig.2), allowing quantitative comparison of their impacts. An additional advantage of this system is that all transgenes are selectively expressed in well-defined neurons (Or47b olfactory sensory neurons) offering exemplary segregation of neuronal compartments facilitating assessment of Tau localization and toxic effects in the soma, dendrites, axons and synaptic fields (Supplementary Fig.1). Or47b neuronal soma and dendrites are in the proximal region of the antennae and their axons innervate and terminate in a specific antennal glomerulus bilaterally, where they synapse with projection neurons relaying olfactory information to higher order learning and memory centres (Supplementary Fig.1). To identify neurons expressing the *Or47b-*GAL4 driver, the membrane tagging *UAS-CD8::EGFP* transgene was recombined onto the same chromosome. Thus neurons expressing the novel Tau transgenes under this driver will be marked by GFP, while Tau expression and its effects will be visualized by tracking the cytoplasmic mCherry (i.e. *mCherry::hTau2N4R)*.

We generated the following Tau mutant transgenic strains: *UAS-hTau2N4R*, the phospho-mimicking mutant *UAS-mCherry::hTau2N4R^E14^*, the aggregation resistant mutant *UAS-mCherry::hTau2N4R^ΔVQIVYK^* (termed hTau2N4R^DEL^), and also a phospho-mimicking but aggregation resistant mutant *UAS-mCherry::hTau2N4R^E14/ΔVQIVYK^*.

Consistent with previous studies, expression of hTau2N4R led to progressive and quantifiable reduction in soma diameter (Fig.1.A7 *versus* Fig.1.A6, C) and an increase in the density of membrane swellings in axon tracts and in axon termini (Fig.2.A2,A2’ *versus* Fig.2.A1, A1’, B), compared to controls. These somatodendritic defects were exacerbated in neurons expressing the phospho-mimicking hTau2N4R^E14^. Degeneration of Or47b neurons was significantly enhanced because even by day 3 (T3), 23% of them and 36% by day 14 (T14) were eliminated compared to hTau2N4R (Figs.1A9, 1B). Moreover, soma diameter of hTau2N4R^E14^-expressing neurons was also significantly reduced by T3 and remained at this level at T14 compared to No Tau and hTau2N4R controls (Figs.1.A9, C). Finally, there was a significant increase in axonal swellings by T3, which became even more prominent by T14 (Fig.2.A4, A4’, B). These data align with other studies indicating that Tau-mediated degeneration is driven by phosphorylation at sites known to be phosphorylated in Tauopathies validating our novel experimental system.

**Figure 1.**
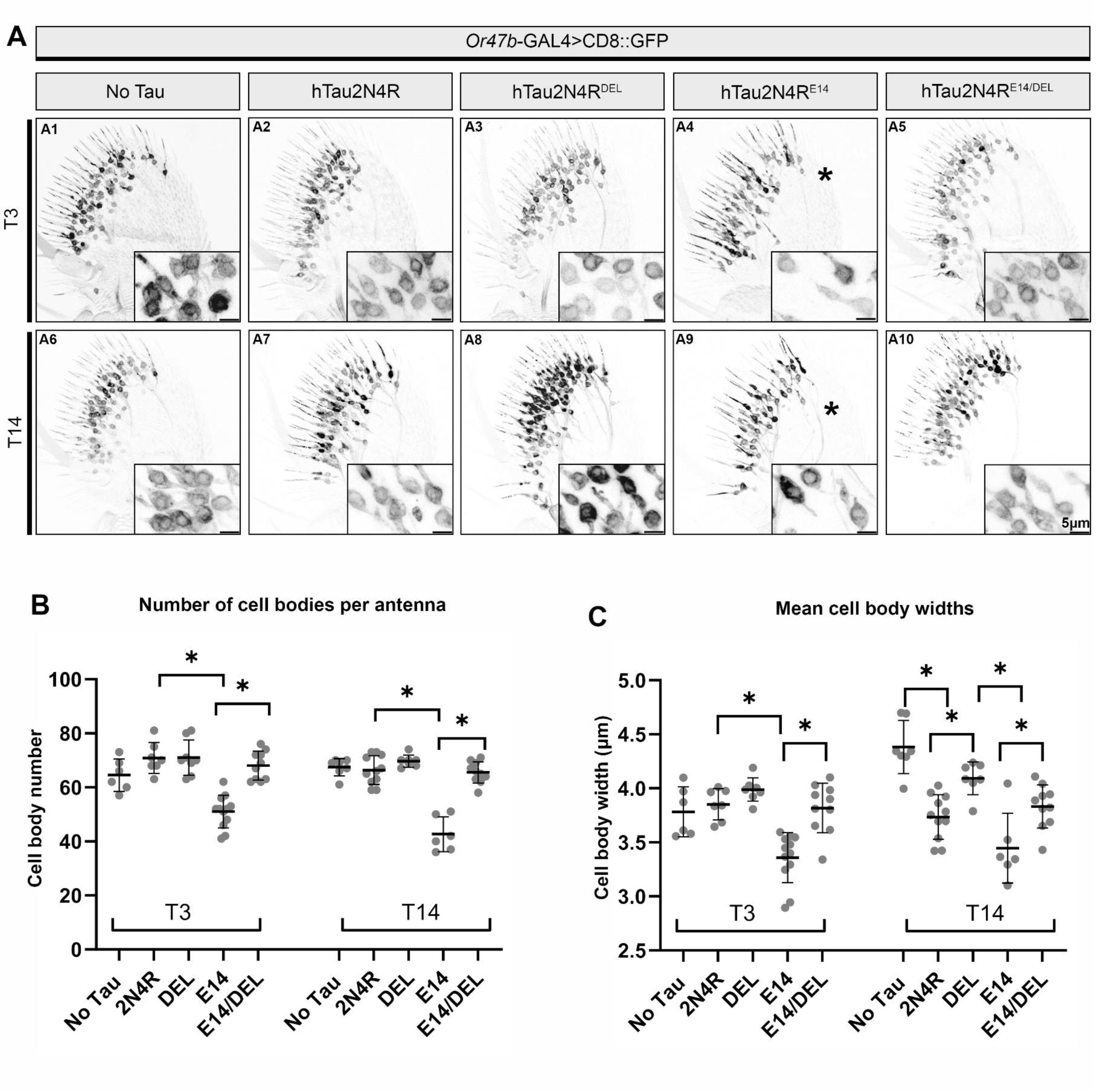
Phosphorylation dependent somatodendritic degeneration requires aggregation. (**A**) Maximum projection confocal images of the adult antenna of flies expressing membrane bound GFP (CD8::GFP) at the Or47b neurons. Columns represent genotypes expressing different mCherry-tagged hTau2N4R variants plus the driver GFP only control (left most column). Rows represent post-developmental age (3 or 14 days). Magnified projections (30 slice depth) of the soma region are included. Asterisk shows the reduced soma density induced by hTau2N4R^E14^ expression (**A9**), which is rescued in hTau2N4R^E14/DEL^ (**A10**). (**B-C**) Quantification of cell bodies per antenna (**B**) and mean cell body width (**C**) for the indicated genotypes and time points. Cell bodies were manually counted and measured using ImageJ (see methods for parameters). Graphs represent Mean ±SD, and dots represent averages per antenna. * = p<0.05 (2-way ANOVA with Tukey’s multiple comparisons). N= 6-12 antennas collected from two biological repeats per genotype.

**Figure 2.**
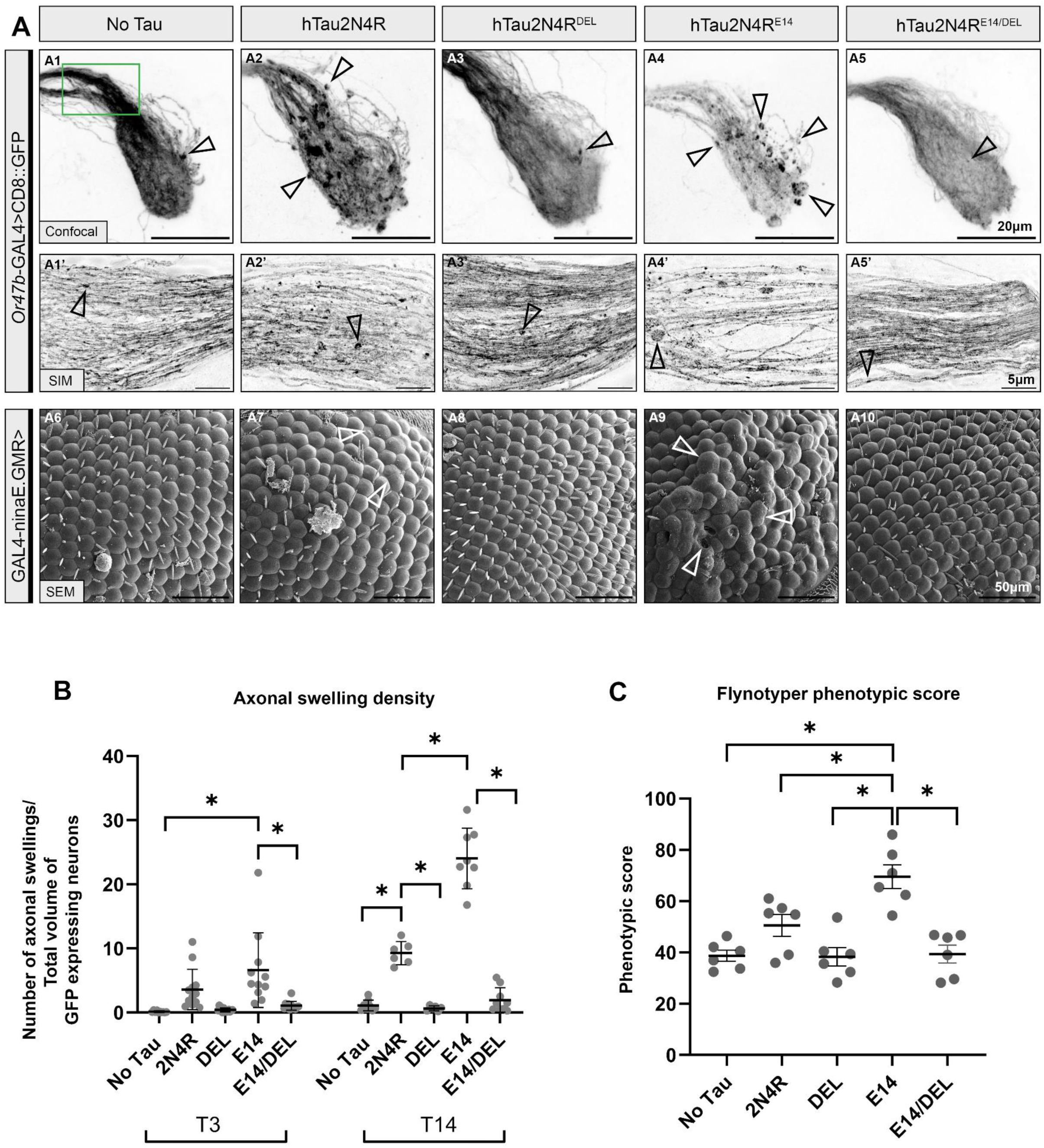
Phosphorylation dependent axonal and synaptic degeneration requires aggregation. (**A**) Maximum projection confocal (**A1-5**) or SIM (**A1’-5’**) images of the brain and antennal lobe of flies expressing membrane bound GFP (CD8::GFP) at the Or47b neurons enhanced with α-GFP. Columns represent genotypes expressing different mCherry-tagged hTau2N4R variants plus the driver GFP only control (left most column). Rows represent post-developmental age 14 days. Normal structure of the neurons shows some axonal swellings (Arrows) that occur with age and are increased in WT hTau2N4R flies (**A2**) and exacerbated in hTau2N4R^E14^ expressing flies (**A4**). Green inset box on B6 highlights the Or47b glomeruli commissure, imaged with SIM (**A1’-5’)** in the middle row. Bottom row of images show SEM micrographs **(A6-A10)** of day 14 *Drosophila* eyes expressing hTau2N4R mutants at 1600x magnification. Using the GAL4-ninaE.GMR driver. Abnormal retina and degenerated cells are labelled with arrowheads. (**B**) Quantification of axonal swellings were automatically quantified using 3D images with IMARIS (see methods). Graphs represent Mean ±SD, and dots represent individual values. * = p<0.05 (2-way ANOVA with Tukey’s multiple comparisons). N= 6-14 brains collected from two biological repeats per genotype. (**C**) Quantification of retinal degeneration using a phenotypic scoring system Flynotyper plugin. Graphs represent Mean ±SD, and dots represent individual values. * = p<0.05 (2-way ANOVA with Tukey’s multiple comparisons). N=6 per genotype.

As aggregation is also implicated in Tau-mediated degeneration and is believed to be triggered by misfolding induced by Tau hyperphosphorylation, we next sought to investigate the impact of reducing the aggregation propensity of toxic potential of hTau2N4R and of hTau2N4R^E14^.

First, we tested hTau2N4R^DEL^ in which the ^306^VQIVYK^311^ domain was deleted. Cell body diameter at T14 was within the control range in these hTau2N4R^DEL^-expressing neurons (Fig.1.A8,C). Similarly, the increase in axon swellings induced by hTau2N4R at T14 was absent in hTau2N4R^DEL^-expressing Or47b neurons (Fig.2.A3. A3’, B). Therefore, deletion of the aggregation-mediating ^306^VQIVYK^311^ motif rendered the toxic hTau2N4R protein benign. Significantly and in striking contrast to the impact of hTau2N4R^E14^, deletion of ^306^VQIVYK^311^ in hTau2N4R^E14/DEL^ completely suppressed the toxic potential of the phospho-mimicking variant. No apparent neuronal death was detectable up to T14 (Fig.1.A10, B) in the hTau2N4R^E14/DEL^ animals and the soma appeared morphologically similar to age matched controls and unlike the sparse and distorted appearance of hTau2N4R^E14^-expressing neurons (Fig.1:A4, A9). Furthermore, cell body diameter was not different in hTau2N4R^E14/DEL^-expressing flies from that of controls at T3 and was similar to that of hTau2N4R-expressing neurons by T14 (Fig.1C). Finally, axon swellings were not apparent either at T3 or T14 (Fig.1.A5, A5’, B). These data indicate that in Or47b neurons, ^306^VQIVYK^311^ deletion renders hTau2N4R non-toxic, even when it is hyperphosphorylated at established pathology-linked sites. Therefore, toxicity of hyperphosphorylated hTau2N4R appears to be mediated by the ^306^VQIVYK^311^ aggregation promoting motif, at least in Or47b neurons.

To test the generality of this remarkable rescue of phospho-Tau-mediated toxicity by deletion of the ^306^VQIVYK^311^ domain, the same Tau variants were expressed in developing photoreceptors, using a *GMR-GAL4* photoreceptor driver. Tau toxicity in photoreceptors manifests as fusion, loss and disorganisation of ommatidia and sensory bristles (Supplementary Fig.3), which can be assigned a ‘disorganization’ phenotypic score using Flynotyper(58)(Fig.2C). Expression of hTau2N4R led to a subtle eye disorganization (Fig.2.A7), which was significantly enhanced by hTau2N4R^E14^ (Fig.2.A9) and quantified (Fig.2C). Notably, these pathologies were totally eliminated upon expression of either hTau2N4R^DEL^ or hTau2N4R^E14/DEL^ (Fig.2:A8, A10). Collectively, these data indicate that toxicity mediated by hyperphosphorylation of hTau is gated through the aggregation-promoting ^306^VQIVYK^311^ domain.

Tau accumulation in axons is reported to impair transport(14, 18) resulting in “axonal swellings” potentially leading to fragmentation and neuronal loss in mammalian Tauopathy models(59–61). These morphological characteristics were examined in hTau-expressing Or47b neurons. Control Or47b neurons present with the typical compact highly fasciculated axons up to T14 (Supplementary Fig.4:A1, Fig.2:A1), with some apparently age-dependent swellings and defasciculation later on (Fig.2:A1, A1’). Significantly however, the number, size and total volume of axonal swellings were highly increased upon hTau2N4R expression at all ages assayed (Supplementary Fig.4:A2, Fig.2:A2). These aberrant morphologies were greatly exacerbated by hTau2N4R^E14^ expression (Fig.2:A4, A4’, Supplementary Fig.4:A4, arrowheads), presenting with prominent axonal swellings and fragmentation, rough dystrophic appearance (Fig.2:A4, A4’, arrowheads), apparently leading to substantial axonal and synaptic loss (Fig.2:A4, A4’), and a decrease in the volume of the neuronal fascicle (Supplementary Fig.5). In contrast, these pathological phenotypes were completely absent upon ^306^VQIVYK^311^ deletion either in hTau2N4R^DEL^ (Fig.2:A3, Supplementary Fig.4:A3), or hTau2N4R^E14/DEL^-expressing animals (Fig.2:A5, Supplementary Fig.4:A5), at all ages tested, which were virtually indistinguishable from age-matched No Tau controls (Supplementary Fig.4:A1, Fig.2:A1). Quantification of axonal swellings and neuronal loss further supports these conclusions (Fig.2:C, Supplementary Fig.5).

Additional confirmation of these conclusions was afforded by Structured Illumination Microscopy (SIM) of the commissural tract located at the top of the olfactory glomeruli (Fig.2:A1) to visualise Tau-mediated axonal degeneration at day 14 (Fig.2:A1’-A5’). Clearly, the increased size of axonal swellings and the extent of degeneration induced by hTau2N4R^E14^ (Fig.2:A4’) were attenuated upon ^306^VQIVYK^311^deletion (Fig.2:A5’).

### Dendritic mislocalisation and axonal accumulation of Tau require the ^306^VQIVYK^311^ domain

Another hallmark of Tau pathology is its mislocalisation or mis-sorting from axonal(62, 63) to the somatodendritic compartments, where it accumulates in large insoluble cytoplasmic hyperphosphorylated inclusions(13, 62–64). Accordingly, at T3 and T14, hTau2N4R and all mutant forms localised throughout the soma, axons and dendrites (Fig.3.B, 4.A, Supplementary Fig.6). However, in the somatodendritic compartments, hTau2N4R and hTau2N4R^E14^ appeared highly concentrated within the dendrites that comprise the antennal sensilla (Fig.3:A1-A10), indicating profound mislocalisation to this compartment. Within the dendrites hTau2N4R distribution became uneven and fragmented from T3 to T14 (Fig.3:A2, A7, Supplementary Fig.6:A2, A7). By comparison, dendritic mislocalisation was greatly exacerbated in an age-dependent manner in hTau2N4R^E14^-expressing neurons (Fig.3:A9, C, Supplementary Fig.6:A4, A9). However, deletion of ^306^VQIVYK^311^ either in hTau2N4R or hTau2N4R^E14^ led to a profound reduction of this mislocalisation, as smaller and sparser Tau accumulates were observed within dendrites (Fig.3:A5, A10 and Fig.3C).

**Figure 3.**
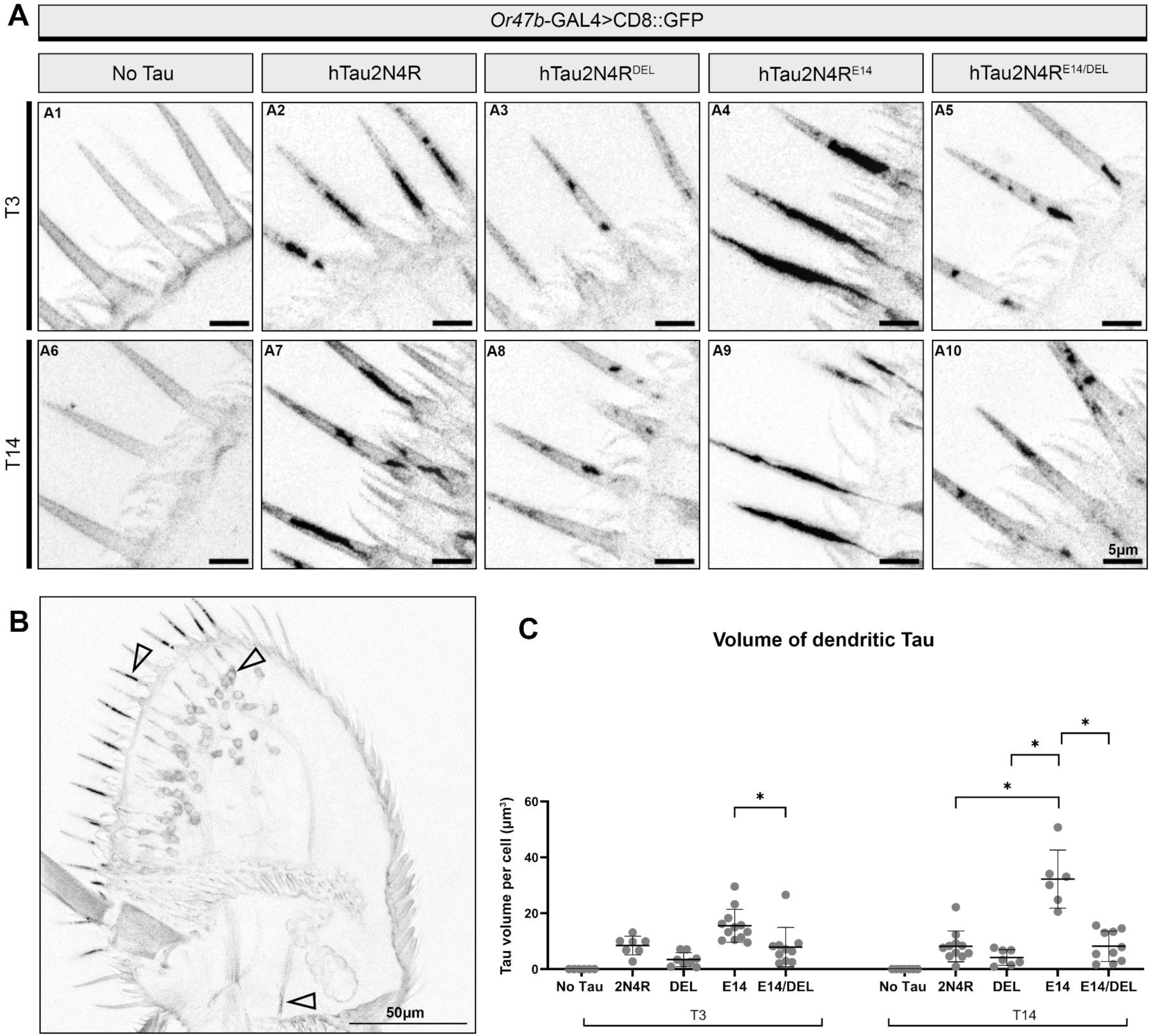
Phosphorylation dependent dendritic Tau accumulation requires aggregation. (**A**) Midline 30 slice stacked confocal images of the adult antenna of flies labelled by the mCherryTau signal from expressed mutants in the Or47b neurons. Columns represent genotypes expressing different mCherry-tagged hTau2N4R variants plus the driver GFP only control (left most column). Rows represent post-developmental age (3 or 14 days). Magnified regions of interest over the dendrites are presented (**A1-A10**). (**B**) Example image of whole mount antenna, image of **A2**. Arrows point toward the Tau accumulation deposited within the dendrites, somas and axons. Other whole mounts can be viewed in supplementary figure 6. (**C**) Quantification of the volume of dendritic Tau in the antenna was quantified using 3D projection images with IMARIS with a surfacing tool using N=6-12 from two biological repeats. Graphs represent Mean ±SD. * = p<0.05 (2-way ANOVA with Tukey’s multiple comparisons).

In the axonal compartment, all hTau protein variants presented with a uniform distribution in Or47b expressing axons and synapses (Supplementary Fig.4:B1-5 of 3 day -old flies. However, by T14 the frequency and size of Tau accumulates (Fig.4:A2) increased in hTau2N4R-expressing flies consistent with the excessive axonal swellings displayed by these axons. These accumulates more than doubled in Tau2N4R^E14^-expressing animals (Fig.4:A4), relative to those accumulating hTau2N4R (Fig.4B). Scrutiny of the axonal bundle using SIM imaging confirms the denser and more prominent intracellular Tau accumulations observed in hTau2N4R^E14^-expressing flies (Fig.4:A4’ *versus* Fig.4:A2’). These pathological Tau accumulates were suppressed by ^306^VQIVYK^311^ deletion, such that by T14 axons were almost devoid of them in hTau2N4R^E14/DEL^ animals. This was confirmed by and SIM analysis demonstrating Tau distribution throughout Or47b neurons, with few accumulates at the synaptic terminals, at the perimeter of the neuronal bundle after T14 (Fig.4:A5, A5’ arrowheads and Fig.4B). Collectivelly then, the aggregation promoting ^306^VQIVYK^311^ domain drives somatodendritic mislocalisation and axonal accumulation of hyperphosphorylated Tau.

**Figure 4.**
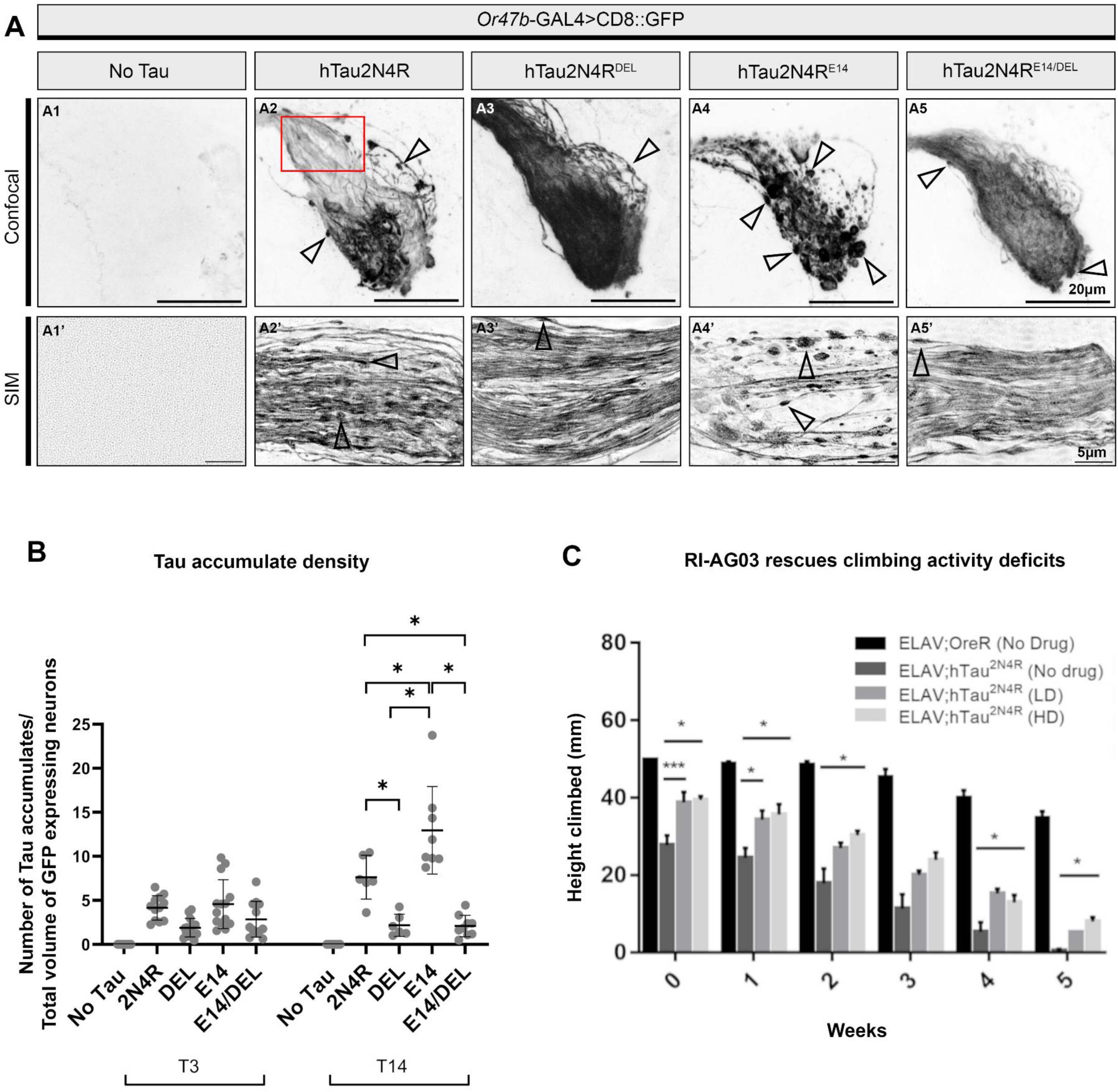
Phosphorylation dependent axonal and synaptic Tau accumulation requires aggregation, and treatment of aggregation inhibitor RI-AG03 rescues locomotive dysfunction in Tau-expressing *Drosophila*. (**A**) Maximum projection confocal (**A1-5**) or SIM (**A1’-5’**) images of the brain, antennal lobe of day 14 old flies expressing mCherry::Tau mutants in the Or47b neurons (enhanced with α-hTau Dako). Arrows point toward beaded Tau accumulates within expressing neurons which increase in frequency and size with age and are exacerbated in hTau2N4R^E14^ expressing flies (**A4**), but are significantly supressed in hTau2N4R^DEL^ (**A3**) and hTau2N4R^E14/DEL^ (**A5**). Red inset box on A6 highlights the Or47b glomeruli commissure, imaged with SIM (**A1’-A5’**) at T14 on the bottom row. N= 6-14. (**B**) Quantification of the Tau accumulate density were quantified using 3D images with IMARIS with a spotting tool (see methods) N=6-14 from two biological repeats. Graphs represent Mean ±SD. * = p<0.05 (2-way ANOVA with Tukey’s multiple comparisons). (**C**) Climbing assay was used to assess locomotive function of panneuronal hTau2N4R expressing flies treated with RI-AG03. Bar graph showing the average height climbed by each group (5 cohorts of 10 male flies) over a 5-week period. Control group (ELAV;OreR) and hTau2N4R-expressing flies were subjected to low-dose (0.08uM) (LD) and high-dose (0.8uM) (HD) treatments of the aggregation inhibitor RI-AG03 from 1 to 5 weeks of age. hTau2N4R-expressing flies exhibited a significant reduction in normal climbing activity compared to control flies. Treatment with RI-AG03 at both LD and HD significantly improved the climbing performance of hTau2N4R-expressing flies. Error bars represent mean ± SEM. * = p<0.0001 (2-way ANOVA with Tukey’s multiple comparisons).

### Peptide inhibition of ^306^VQIVYK^311^ rescues Tau-mediated locomotive deficits

To explore the translational potential of targeting the ^306^VQIVYK^311^ domain, we tested whether targeting it pharmacologically could suppress pathological manifestations. To that end, flies expressing hTau2N4R panneurally were fed a low-dose (0.08μM referred to as LD) and high-dose (0.8μM referred to as HD) of the aggregation inhibitor peptide RI-AG03. This inhibitor is non-toxic to *Drosophila* at these doses, and it effectively blocks the ^306^VQIVYK^311^-mediated aggregation(40). We assessed adult fly climbing efficiency of adult flies in a well-established test to assess the effects of Tau in the fly CNS and peripheral nervous system (PNS)(34, 38). We quantitated the climbing of control flies over 5 weeks, and found a progressive but modest 24% decline in climbing efficiency. In contrast, panneuronal expression of hTau2N4R reduced climbing efficiency by 40% even in 1 day-old flies (T1). By week 3, a 60% decline was observed and by week 5, hTau2N4R-expressing flies were unable to climb at all. In contrast, treatment with RI-AG03 significantly improved climbing of hTau2N4R-expressing flies, though not to control levels (Fig.4C). Although the peptide targets and inhibits the ^306^VQIVYK^311^ domain, its effects are unlikely as potent as deletion of this sequence, as drug efficacy is highly dependent on its stability, penetration, assimilation and pharmacokinetics. Nonetheless, these data suggest that pharmacological inhibition of the ^306^VQIVYK^311^ domain is likely an effective method to suppress hyperphosphorylation and aggregation-dependent Tau phenotypes.

## Discussion

Phosphorylation and aggregation are both established pathogenic post translational modifications of Tau thought to play a central role in mediating Tau-toxicity. They therefore represent significant disease-modifying therapeutic targets for the treatment of Tauopathies. Yet the causal relationship between these distinct post-translational modifications and their relative contribution towards Tau toxicity is not clear. Our study determines the relative contribution of hyperphosphorylation and aggregation in Tau-mediated neurodegeneration in aging *Drosophila* olfactory neurons. We show that ^306^VQIVYK^311^ is an excellent candidate for therapeutic intervention *in vivo*, and provide novel data which shows that its targeting can render even highly phosphorylated Tau species inert.

### Phosphorylation as a Key Driver of Tau-mediated Toxicity

We demonstrate that the pseudo-phosphorylated hTau2N4R^E14^ mutant significantly exacerbates neurodegenerative phenotypes compared to hTau2N4R across various neuronal compartments in a novel experimental system. This finding aligns with previous research using TauE14 and other phospho-mimicking mutants which have consistently shown increased neuronal loss and organizational defects when expressed in alternative *Drosophila* neuron subtypes including the mushroom bodies(65), retinal neurons(20, 66), and circadian clock neurons(67). However, focusing on Or47b neurons revealed not only severe morphological deficits, such as profound loss of somatic and axonal tissue, but also subtler hallmark manifestations more difficult to document *in situ* such as the marked increase in axonal swelling size and frequency.

While TauE14 is an aggressive mutant mimicking extensive hyperphosphorylation akin to that seen in AD, it effectively recapitulates the pathogenesis associated with pathological Tau hyperphosphorylation. This underscores the causal role of the 14 GSK-3β phosphorylation sites mutated in TauE14, in Tau-mediated toxicity and validates the focus on reducing phosphorylation as a therapeutic strategy. Adding to this we now show, for the first time, that the pathogenic potential of these sites requires the aggregation promoting ^306^VQIVYK^311^ domain.

### The Critical Role of the ^306^VQIVYK^311^ Domain in Tau Toxicity

There is significant evidence suggesting that phosphorylation is a precursor for aggregation. Using pseudo-phosphorylated Tau mutants, multiple groups have identified a number of phosphorylation sites that act in concert to promote polymerization of monomeric Tau(44–46, 51, 68–70) and which increase the aggregated sarkosyl-insoluble Tau fraction in *Drosophila,* including TauE14(20, 66).

Importantly, our results reveal that the toxicity driven by hyperphosphorylation is fundamentally dependent on the aggregation-promoting ^306^VQIVYK^311^ domain. Remarkably, deletion of this domain in both hTau2N4R and hTau2N4R^E14^ almost completely eliminated toxicity, despite persistent phosphomimicking at disease-associated epitopes. Consistent with our findings, a previous study similarly showed that deleting the ^306^VQIVYK^311^ motif in a *Drosophila* model expressing hTau0N4R attenuated Tau toxicity, rescuing retinal degeneration and shortened lifespan (38).

Collectively this highlights the therapeutic potential of targeting this domain *in vivo*. Building on this, we provide novel insight into its critical role within the context of disease-related hyperphosphorylation, suggesting that the gain-of-function (GOF) toxicity driven by ^306^VQIVYK^311^-dependent mechanisms surpasses any phosphorylation-mediated loss-of-function (LOF) or GOF effects in contributing to overall Tau toxicity.

### The Interplay Between Phosphorylation and Aggregation

Our data provides compelling evidence that Tau aggregation is downstream of and gates the toxicity of hyperphosphorylated Tau. We show that hTau2N4R^E14^ significantly increases both the number and size of Tau accumulations within axons, consistent with findings in murine hippocampal neurons(68). Several mechanisms may explain how Tau phosphorylation promotes aggregation, including changes in electrostatic charges that create seed-competent species(71, 72), and reduced microtubule binding, both of which can favour aggregation conformation potential(47, 73, 74), and shift the microenvironmental equilibrium towards aggregation.

Importantly, our results demonstrate that the ^306^VQIVYK^311^ domain is crucial for facilitating Tau accumulation and subsequent neuronal degeneration, regardless of the specific mechanism by which phosphorylation enhances Tau aggregation. This is corroborated by previous studies showing that deletion of this motif significantly impedes aggregation and β-sheet assembly of both 3R and 4R Tau proteins *in vitro* and *in vivo*(38, 54, 75).

### Impact on Tau Mislocalisation and Accumulation

The mislocalisation of Tau away from the axonal compartment is well documented in patients with primary Tauopathies(76). Previous studies have shown that the phosphorylation status of Tau alters its motility and localisation, contributing to its somatodendritic localization(13, 77).

Our study reveals that mimicking hyperphosphorylation leads to greater dendritic mislocalisation and accumulation of Tau, while deletion of the aggregation-promoting domain reduces or delays this effect. The increased mislocalisation of phosphorylated Tau may be due to its detachment from microtubules and subsequent transport to the somatodendritic compartment. The fact that aggregation-resistant Tau exhibits reduced dendritic mislocalisation suggests that the process of aggregation plays a role in sequestering Tau within the somatodendritic compartment.

### Implications for Therapeutic Strategies

Disease-modifying treatments for Tauopathies invariably aim to reduce pathogenic Tau species, whether that is through anti-sense oligonucleotide(78) or vaccination(79) mediated clearance, reduction of Tau phosphorylation through kinase inhibition(80–82) or suppression of Tau aggregation through the use of aggregation inhibitors(39, 40, 55, 83, 84).

Our findings have significant implications for the design of Tau-centred disease-modifying therapeutics. While reducing phosphorylation remains a valid approach, our results suggest that targeting the ^306^VQIVYK^311^ domain may be a more effective strategy to suppress the toxicity of multiple pathogenic forms and isoforms of Tau. This domain appears to be a critical point through which various pathogenic mechanisms, including hyperphosphorylation, exert their toxic effects. Indeed several studies have reported promising findings showing the neuroprotective effect of peptides targeting this and the ^306^VQIVYK^311^ domain. We also show that using the peptide aggregation inhibitor RI-AG03(40) to pharmacologically recapitulate the deletion of the ^306^VQIVYK^311^ domain reduces age-related Tau-mediated behavioural deficits in Tau transgenic flies.

Further studies should investigate how deletion of the ^306^VQIVYK^311^ motif affects other cellular processes, such as microtubule binding efficacy, cell signalling, and downstream degradation pathways. Additionally, exploring the effects of peptide aggregation inhibitors on other neurobiological functions of Tau will be crucial for translating these findings into effective therapeutic interventions.

## Conclusion

In conclusion, our study provides compelling evidence that while Tau phosphorylation is a key driver of toxicity, its pathogenic effects are primarily mediated through the aggregation promoting ^306^VQIVYK^311^ domain. This insight offers a novel perspective on the hierarchy of pathogenic events in Tauopathies and that regardless of the level of Tau phosphorylation, if aggregation is inhibited early enough the toxicity of Tau can be reduced.

## Figure Titles and Legends

**Supplementary Figure 1. The Or47b expressing olfactory sensory neurons.**

Panel A. Schematic panel orientation of the adult *Drosophila* brain, showing the location of the olfactory receptor 47b (Or47b) neurons expressing membrane bound GFP (green), co-expressing mCherry tagged human Tau2N4R (mCh::hTau) (magenta). Representative confocal images of whole mount (**A**) Antenna and (**C**) the Antennal lobe (AL) is shown. (**B**) Schematic diagram of the Or47b olfactory neurons. The AL is outlined in dashed black, containing multiple olfactory glomeruli, including Or47b (green) containing a bundle of ∼60 olfactory receptor neurons (ORNs). ORN cell bodies are contained within the 3^rd^ antennal segment of the Antenna (**A, A’)** and its dendrite projects into the sensilla hairs of the antenna. The axons project from the antenna connecting into the (**C**) AL of the brain, containing the (**C’**) axonal bundles and synaptic terminals of the Or47b neuron. For simplicity, only a few of the 60 ORNs are shown. Made using Biorender.

**Supplementary Figure 2. Tau is equally expressed by all mutants.**

**(A)** Western immunoblot analysis of total Tau (α-hTau Dako), taken from 15 male, 2 day old homogenised transgenic fly heads expressing wildtype mCherry::hTau2N4R and mutant forms. Driver is GMR.ninaE-GAL4. Representative blot is shown (**A**) and mean quantification plotted as a bar graph (**A’**) (n=3 biological repeats). Error bars presented as means with SEM. Actin was used as the loading control. mCherry::hTau constructs each run at ∼100kDa due to the extra weight of the fluorescent tag (hTau2N4R usually runs at ∼70kDa) and Actin runs between 35 and 55kDa, at ∼40kDa.

**Supplementary Figure 3. *Drosophila* Eye diagram, phosphorylation dependent retinal degeneration requires aggregation.**

**(A)** SEM micrograph of whole *Drosophila* head. White insert shows the posterior region of interest where Tau-mediated retinal degeneration occurs using the GAL4-ninaE.GMR driver. (**A’**) The *Drosophila* eye comprises of ∼800 single ommatidia units (highlighted in yellow) which are morphologically sensitive to toxic proteins like Tau. Loss of sensory bristles (blue arrows) are also markers of retinal health, which are viewed using SEM techniques.

**(B)** Digital microscope images (600x) of *Drosophila* eyes expressing each hTau2N4R mutant. A rough appearance of the posterior half of the eye appears after hTau2N4R expression (**B2)** and is exacerbated by the TauE14 mutations (**B4).**

**Supplementary Figure 4. T3 Axonal degeneration and Tau accumulation.**

**(A)** Maximum projection confocal images of the adult antenna of flies expressing membrane bound GFP (CD8::GFP) at the Or47b neurons. Columns represent genotypes expressing different mCherry-tagged hTau2N4R variants plus the driver GFP only control (left most column).

**(B)** Maximum projection confocal (**B1-5**) images of the brain, antennal lobe of day 3 old flies expressing mCherry::Tau mutants in the Or47b neurons (enhanced with α-hTau Dako). Arrows point toward beaded Tau accumulates.

**Supplementary Figure 5. Total volume of GFP expressing neurons.**

Quantification of neuron volume was automatically quantified using 3D images with IMARIS (see methods). Graphs represent Mean ±SD, and dots represent individual values. * = p<0.05 (2-way ANOVA with Tukey’s multiple comparisons). N= 6-14 brains collected from two biological repeats per genotype.

**Supplementary Figure 6. T3 and T14 Somatodendritic Tau accumulation in whole mount antenna.**

Midline 30 slice stacked confocal images of the adult antenna of flies labelled by the mCherryTau signal from expressed mutants in the Or47b neurons. Columns represent genotypes expressing different mCherry-tagged hTau2N4R variants plus the driver GFP only control (left most column). Rows represent post-developmental age (3 or 14 days).

**Supplementary Figure 7. Using IMARIS software for quantification of Tau accumulation and neuronal degeneration using “spots” and “surfaces” masking tools.**

## Methods and materials

### Fly stocks

*Drosophila melanogaster* were grown on standard cornmeal media in a controlled 12/12 hour light/dark cycle at constant humidity. Each genetic cross was set at 18°C, and selected progeny was collected and moved to 29°C to age for 3 or 14 days. The UAS/GAL4 binary expression system was used for human Tau expression(85) (85), with drivers Or47b-GAL4 and ninaE.GMR-GAL4.

### Transgenic fly lines

#### GAL4 driver lines

Olfactory driver +/+ OR47b-GAL4,UAS-mCD8::GFP/Cyo;+/+ for expression of Tau variants in a small bundle of olfactory sensory neurons (Odorant receptor 47b – VA1d glomeruli) was used for confocal and structured illumination microscopy. Retinal driver +/+; GAL4-ninaE.GMR/Cyo; +/+ for strong expression of Tau variants in photoreceptor cells, utilised for western blot immunoassays.

#### Tau lines

All UAS-Tau lines (UAS-mCherry::hTau2N4R, UAS-mCherry::hTau2N4R^Δ306VQIVYK311^, UAS-mCherry::hTau2N4R^E14^ and UAS-mCherry::hTau2N4R^E14-Δ306VQIVYK311^) were created by the Allan Lab (University of British Colombia, Canada). Tau E14 is generated by mutating 14 AD-relevant epitopes Thr111, Thr153, Thr175, Thr181, Ser199, Ser202, Thr205, Thr212, Thr217, Thr231, Ser235, Ser396, Ser404 and Ser422 to glutamate(13). All UAS-Tau variants were integrated into the same attp40 site on the second chromosome, ensuring the same expression of each Tau variant to allow for direct comparison of toxicity (Supplementary Figure 2).

### Confocal imaging experimental procedure

#### Tissue preparation

##### Drosophila brains

This study utilised the *Or47b*-GAL4 driver. Adults were collected 0-1 days after eclosion and moved from 18°C →29°C to be aged for either 3 or 14 days. Flies were anesthetised with CO2 and the males were selected for dissection. Heads were removed and fixed for 20 minutes in 4% paraformaldehyde (PFA) in phosphate-buffered-saline with 0.2% Triton-X 100 (PBST). The heads were then washed 3 times, for 20 minutes each in PBST, and then dissected under a light microscope in PBST. Brains were then blocked in 3% Normal Goat Serum in PBST (LAMPIRE Biological Laboratories) for 1 hour. Then, primary antibodies were added and incubated overnight. The following day, the primary antibodies were removed and the brains were washed 3 times, for 10 minutes each in PBST. The brains are then left to incubate with the secondary antibodies for 2 hours. Before mounting, the brains are again washed 3 times, for 10 minutes each in PBST. Primary antibodies used: chicken-Anti-GFP (1:1000)(Abcam ab13970), α-human Tau (1:1000) (rabbit-Anti-human-Tau Dako). Secondary antibodies used: Goat-Anti-Rabbit-Alexa-555 (1:500) (Abcam Ab150078), Anti-Mouse-Alexa-488 (1:500) (Invitrogen A11039). ***For confocal imaging*** the brains are then transferred to a poly-L-Lysine coated slide containing coverslip well, and mounted using Vectashield® hard-set mounting media (H-1400-10). Slides were stored in a freezer (−20°C) and imaged within 3 days. ***For axonal and synaptic structured illumination microscopy (SIM) imaging***, after antibody incubation brains were post-fixed in 4% PFA for 10 minutes, washed and mounted on Superfrost plus adhesion glass slides. Prolong Gold^TM^ is placed on top of the brains and a high-precision thin coverslip (18×18 mm 170± 5µm gently layered on top. Slides left to cure for 24 hours (at room temperature in the dark) before being imaged.

##### Drosophila antenna

The same progeny was used for antennal dissection. Only one antenna was taken from each animal to avoid the same repeats from the same animal. One antenna was removed from each animal and fixed for 20 minutes in 4% PFA with 0.2% PBST followed by 3 PBST washes and immediate mounting onto a poly-L-Lysine coated slide using Vectashield hard-set mounting media. No antibodies were used because the adult antennal cuticle is too thick and highly autofluorescent, instead the endogenous mCherry and CD8::GFP fluorophores were captured during imaging.

##### SEM, Drosophila eye

14 day old flies were euthanised, fixed in 4% formaldehyde 3% glutaraldehyde in 0.1M PIPES buffer. Samples were rinsed in 0.1M PIPES buffer twice, immersed in 2% osmium tetroxide for 1 hour, then dehydrated for 20-minutes at each ethanol concentration: 30%, 50%, 70%, 80%, 90%, 100%, 100%. Samples were critical point dried, mounted on aluminium stubs before sputter coating with gold/palladium in a Polaron E5100 sputter coater. Samples were examined using a Quanta 250 FEG-SEM.

#### Image acquisition

##### Confocal Image acquisition

Images of the entire *Drosophila* brain and antenna were taken with a Leica SP8 line scanning confocal microscope. Images were taken with 12-bit depth stacks at 63x magnification in glycerol immersion oil, taken at 1024×1024 resolution. 488nm laser was used to detect degeneration of the Or47b neurons and 555nm laser to detect Tau signal. Two sequential excitation steps are used to avoid bleed through between channels. Representative images shown are presented as full Z-stacks, unless otherwise stated. As only structural parameters were evaluated, confocal acquisition settings were adjusted if necessary to capture all structural information.

##### SIM image acquisition

Commissure tract of the Or47b axons within the brain were imaged via SIM (DeltaVision OMX Flex super-resolution microscope) with a 63x Immersion oil objective. An immersion oil with a refractive index of 1.510 (Cargille Laboratories USA) was used whilst objective collar settings were adjusted to 013 to provide high-resolution images.

#### Image and statistical analysis

Quantitative analysis of axonal swelling density, Tau accumulate density, volume of dendritic Tau per cell and total volume of GFP expressing neurons were performed using the IMARIS (Supplementary Fig.7) 3D image analysis software. Quantification of cell body density and cell body widths was performed in ImageJ using the cell counter tool and measuring tool respectively. Only one antenna was imaged from one fly. Both hemispheres of the brain were analysed, however in some instances where only one hemisphere was available, quantification was doubled.

Quantification was always carried out using defined parameters. To quantify the number of axonal swellings, the IMARIS “spotting” tool was used to discretely measure the number of axonal swelling surfaces of ≥1.2 µm with a surface detail of 0.36 μm. Axonal swelling density was then calculated using the total volume of GFP expressing neurons, which was measured using the “surfacing” tool detecting structures ≥0.5 µm and a surface detail of 0.2 µm. The number of Tau accumulates in the brain were defined using the “spotting” tool for accumulates that had a diameter between 1.5-2.5 μm, with a surface detail of 0.36 μm. The volume of dendritic Tau accumulates was measured using the “surfacing” tool detecting accumulates ≥1.0 μm defined by surface detail of 0.36 μm. Thresholding was achieved with local contrast background subtraction with the diameter of the largest sphere which fits into the object. Thresholding was manually adjusted for each animal to capture true structures. Analysis of each cohort was completed under blinded conditions. Statistical analysis was performed using a Two-way ANOVA and a multiple comparisons test.

Quantitative analysis of retinal degeneration was achieved using digital microscope images established ImageJ plugin tool Flynotyper(58) n=6.

#### SDS-PAGE Western blotting procedure

##### Tissue preparation

This study utilised the GMR.ninaE-Gal4 driver. Flies were collected at 18°C and moved to 29°C to age for 2 days for total Tau expression and 14 days for measuring p-Tau levels. Flies were then snap frozen in liquid nitrogen and decapitated, in preparation for homogenisation. The heads and brains were then homogenised in an Eppendorf using a pestle for 3 minutes in ice-cold fly homogenising buffer (FHB) containing 10mM Tris, 150mM NaCl, 5mM EDTA, 50mM NaF, 1mM Na_3_VOF_3_, 5mM NaPPi, 4mM Urea, 5mM DDT, Glycerol, Protease inhibitor (1:100) and Phosphatase inhibitor (1:100). 3ul of FHB was used per head. Lysates were then centrifuged at 10,000 x *g* for 5 mins at room temperature. Once centrifuged, the supernatant was collected and boiled with 2X SDS-PAGE sample buffer in a 1:1 ratio at 95°C for 5 minutes in a sterile Eppendorf.

##### Western blots

Proteins were resolved using vertical electrophoresis on 1.5mm 10% SDS-PAGE acrylamide hand-cast gels at 120V for 90 minutes.

##### For total Tau expression

15 male flies were collected for each set of repeats for each sample group. Each well used contained 25µL of the fly homogenate sample buffer mix. Page Ruler Protein Plus ladder (Thermofisher Scientific) was added to each gel. After electrophoresis, the gel containing resolved proteins were electro-transferred onto a nitrocellulose membrane using the electroporator at 60V for 90 minutes, with an icebox to maintain ∼4°C. Following electro-transfer, the membranes were immediately placed in blocking buffer containing 5% bovine serum albumin (BSA), 0.1% tween using 1X PBS on a shaker for 60 minutes. Following blocking, the membranes were then incubated with the primary antibodies in a cold room shaker maintaining 4°C overnight. Antibodies listed in table 1 were made up in the blocking buffer at the given concentrations. Actin was the loading control used in all experiments. The membranes were then washed 3 times with 1X PBS 0.5% tween for 5 minutes on a shaker. Lastly, the membrane was incubated in Anti-Rabbit-800 (Licor, #D00804) and Anti-Mouse-680 (Life technologies A21057) (1:20,000), and washed 3 times with 1X PBS 0.5% tween, for 5 minutes. The membranes were then visualised using LiCOR imager, and analysed using ImageJ software.

**Table 1:**
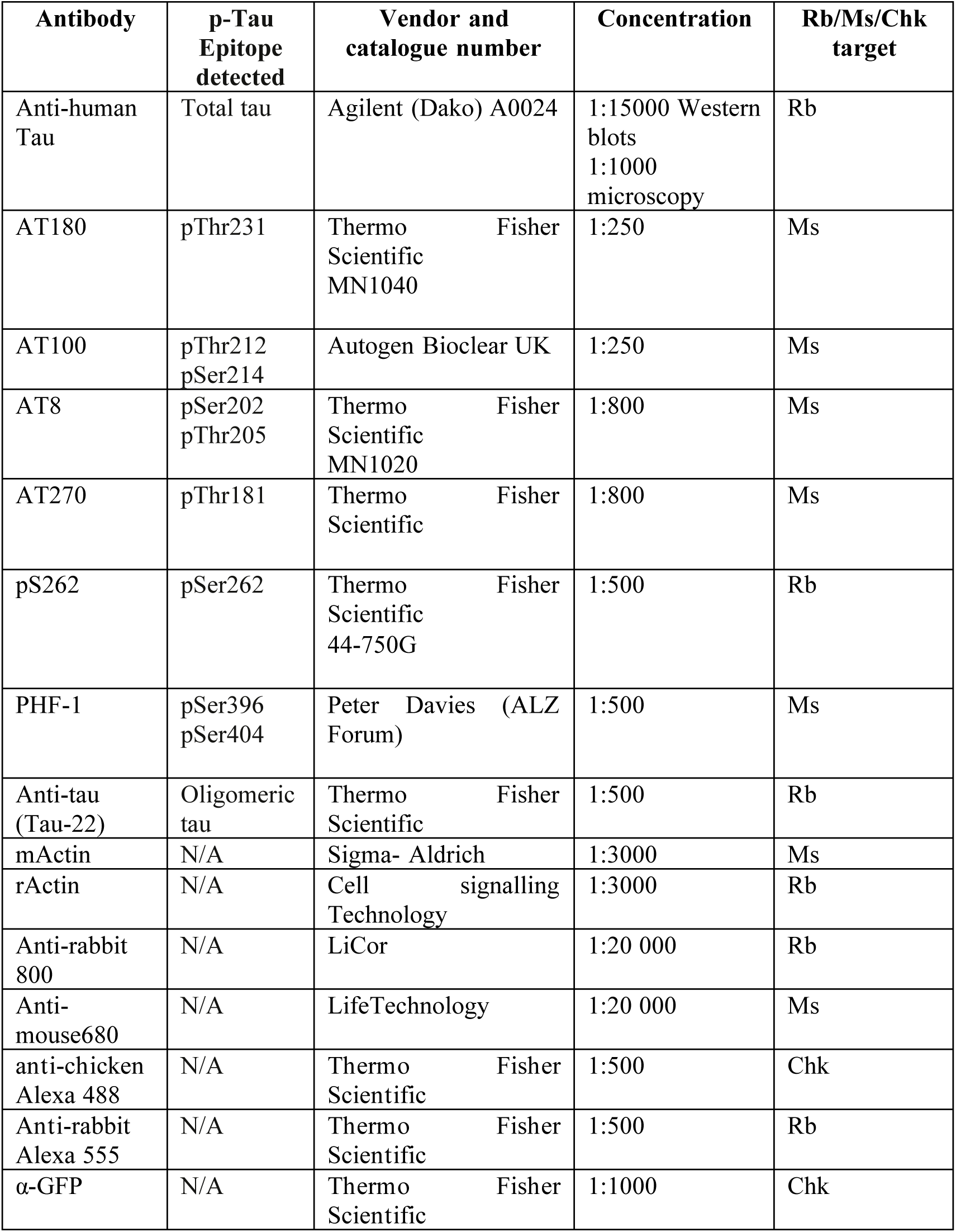
Table of antibodies used in immunolabeling for microscopy, and western blot immunodetection. Further antibody classification as either rabbit/polyclonal (Rb) or mouse/monoclonal (Ms). Phosphoepitopes detected at serine (Ser) or threonine (Thr) residues.

**Table 2:**
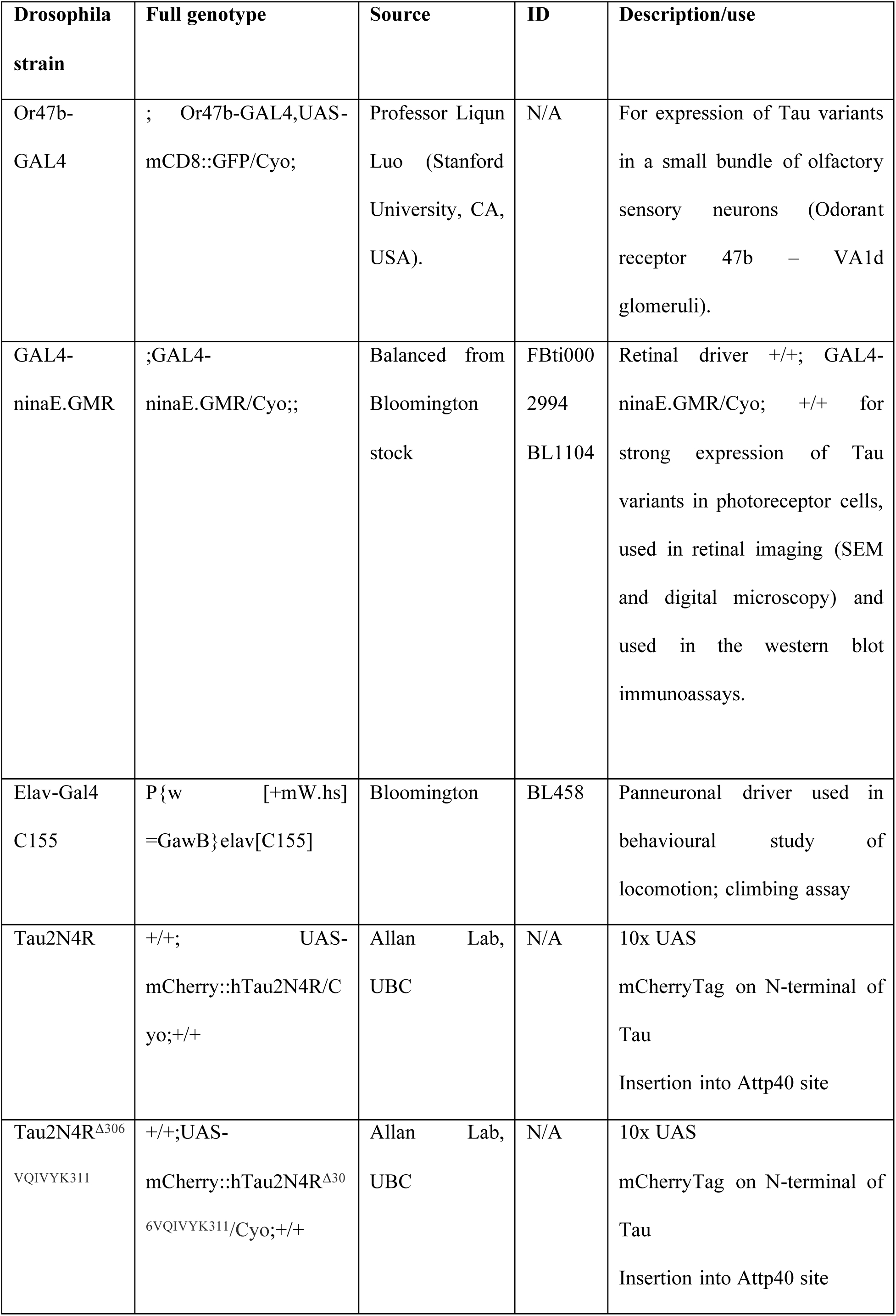
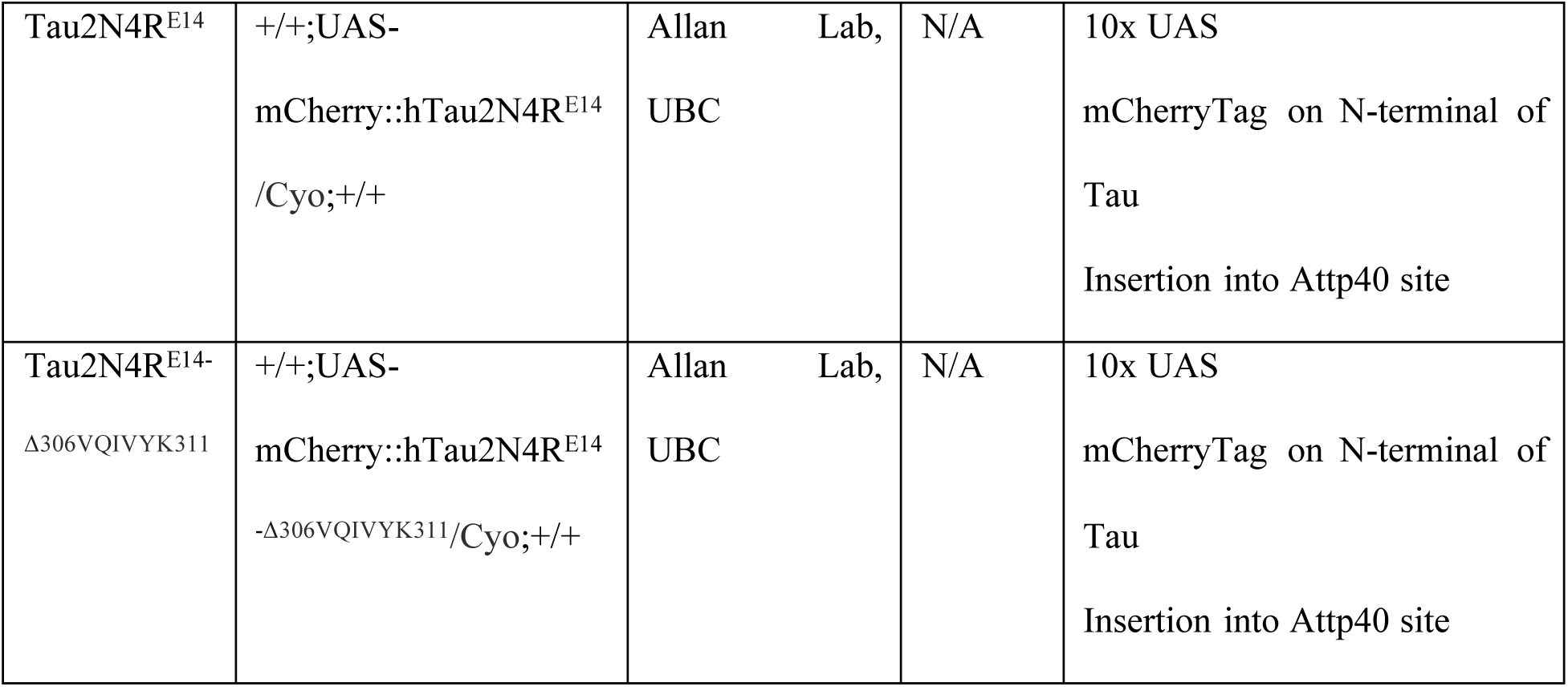
Table of *Drosophila* stocks, genotypes, and source.

##### Statistical Analysis

Western blot bands were quantified using ImageJ software. Tau bands were measured at ∼100kDa and Actin bands were measured ∼40kDa.

##### Climbing assay

Climbing behaviour was performed using 5 cohorts of 10 adult male flies (n= 50) with panneuronal expression (*Elav*-GAL4) of hTau2N4R and Oregon R WT as a control (WT). Flies were placed in cylinders in an assay room with controlled lighting conditions, temperature (23 °C) and humidity (30–40%). The assay was conducted by tapping flies 3 times upon a mouse pad to send the flies to the bottom of the cylinder and recording the flies climbing up the sides of the measuring cylinder. The distance climbed by the flies was recorded at 10 seconds. The assay was repeated three times, with 2 minute rest between each trial and the mean was calculated. Crosses were set on RI-AG03 enriched diet, and progeny were aged on the drug throughout the experimental period at either a low (LD) and high (HD) dose of the RI-AG03 tau aggregation inhibitor (0.08 µM and 0.8 µM). These concentrations were selected after preliminary studies (40). 2-way ANOVA was used for Statistical Analysis via Graphpad Prism Software.

## Notes

### Competing Interest Statement

The authors have declared no competing interest.

