## Supplementary figures for "Aggregation promoting sequences rather than phosphorylation are essential for Tau-mediated toxicity in *Drosophila*"

**Supplementary Figure 1**

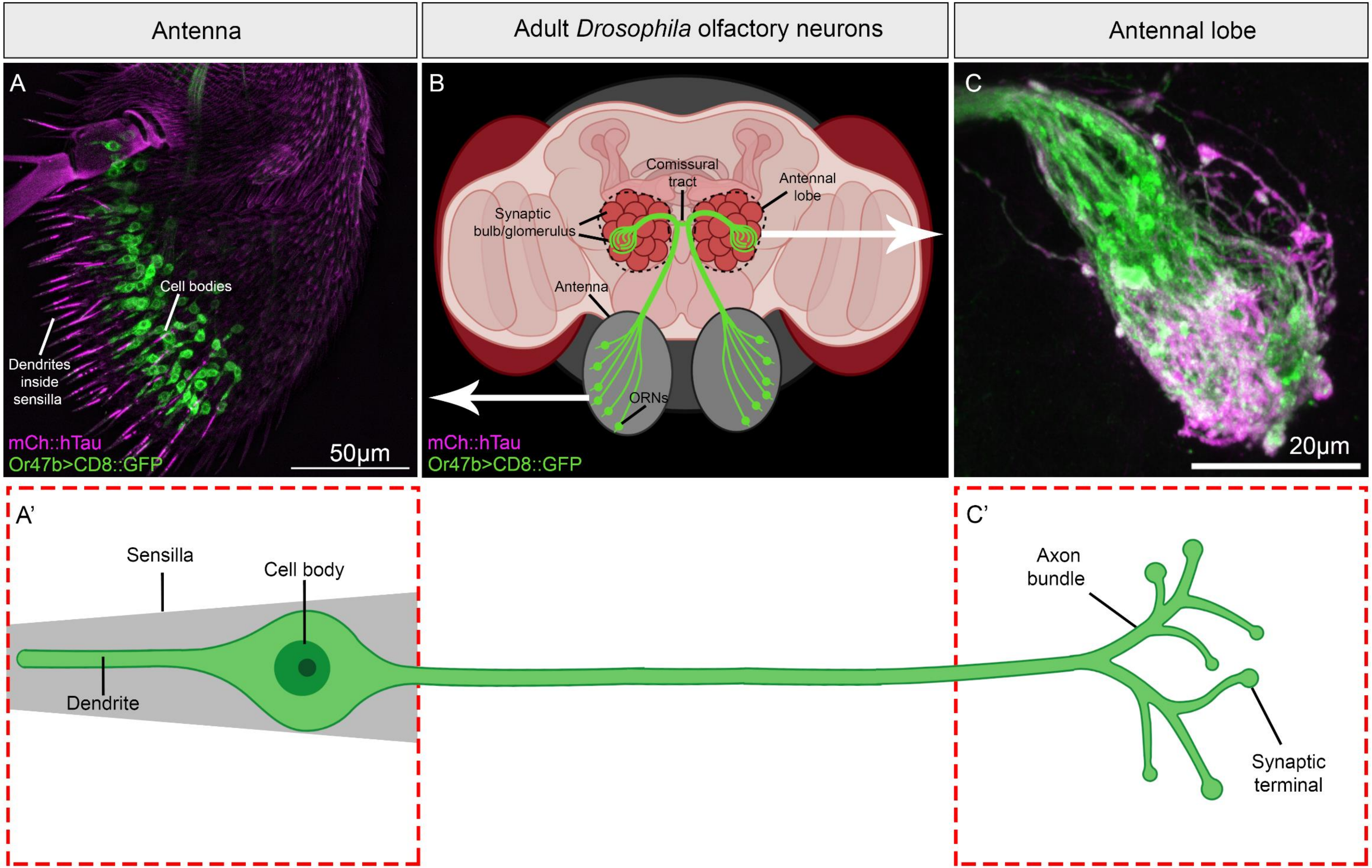

**Supplementary Figure 2**

A

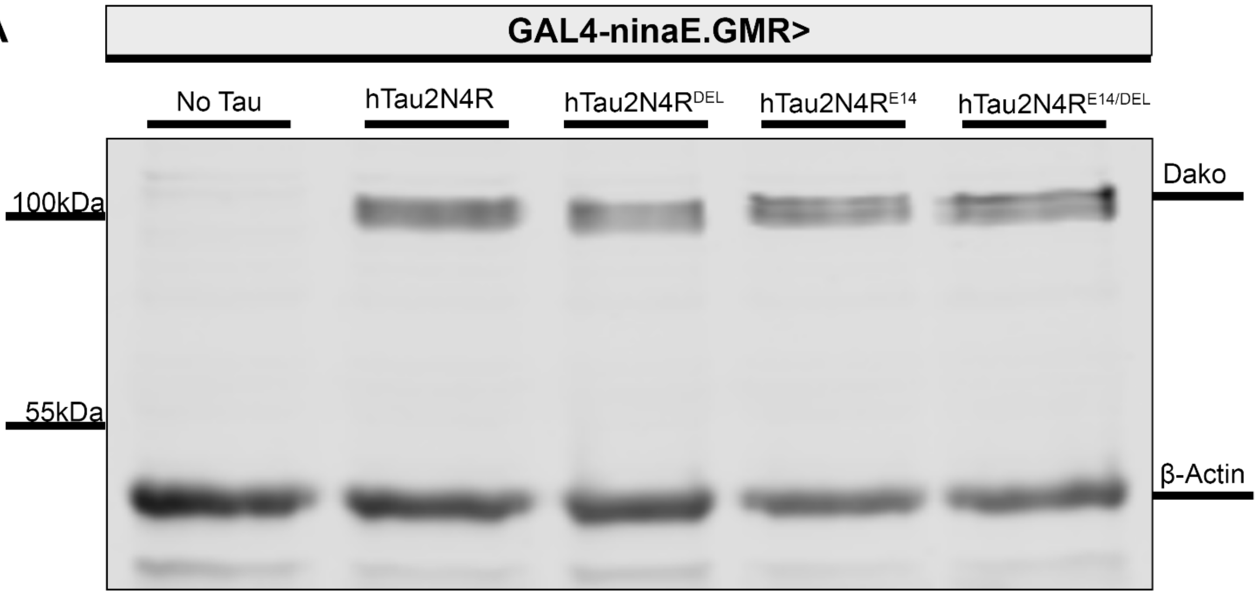


A’

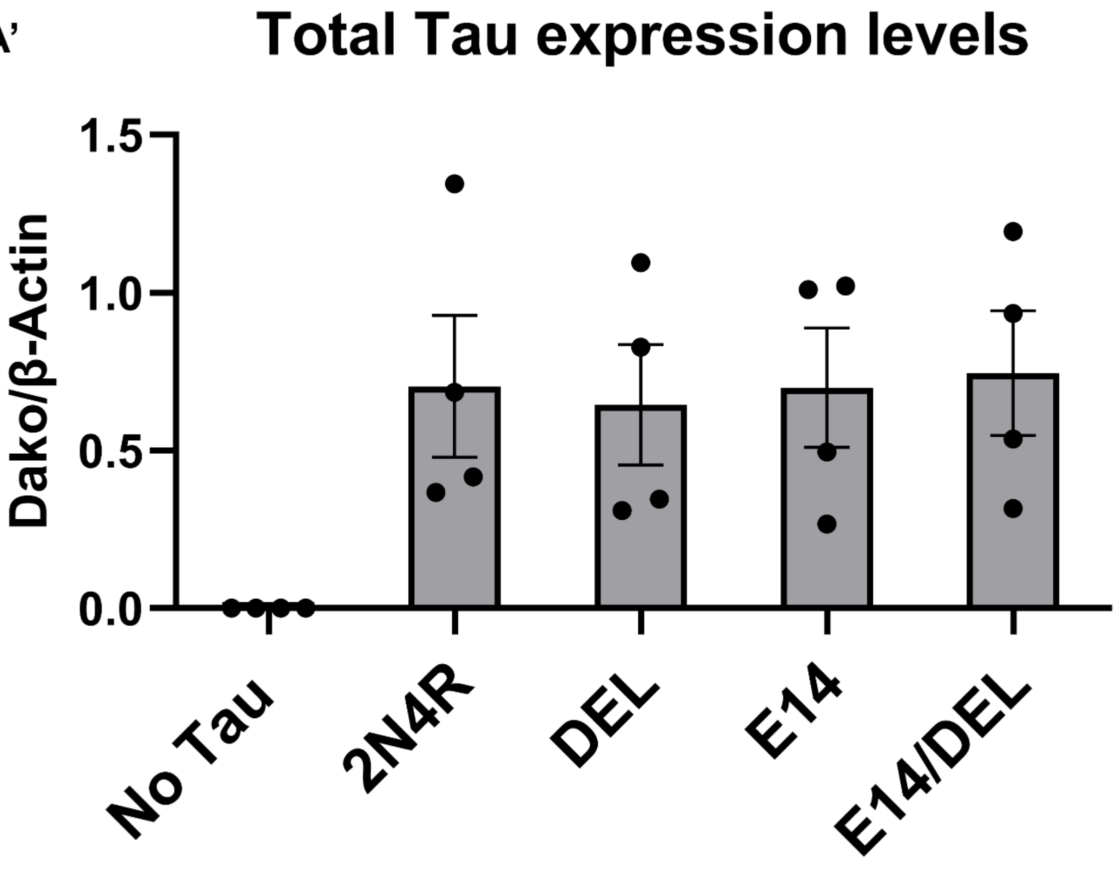

**Supplementary Figure 3**

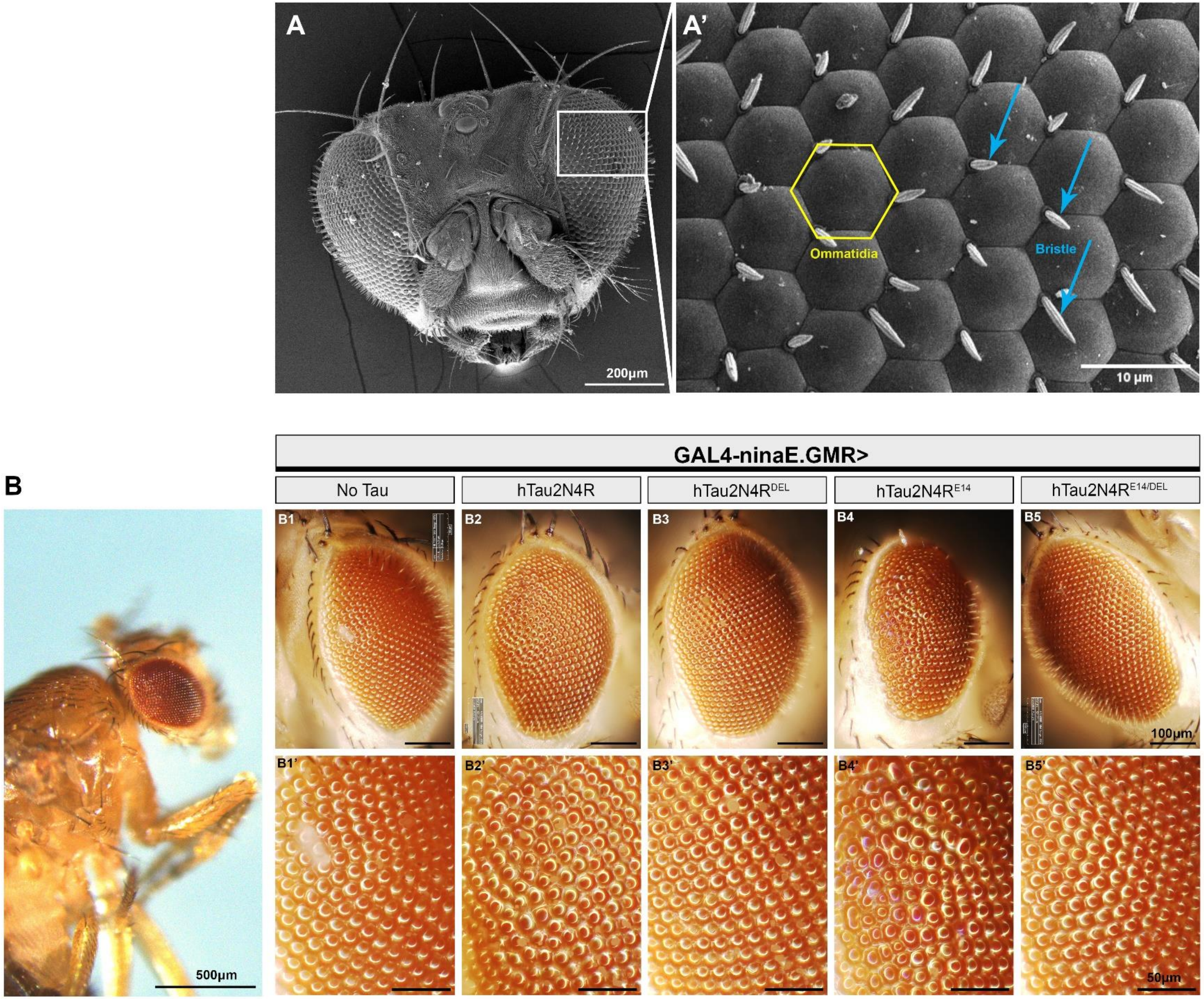

**Supplementary Figure 4**

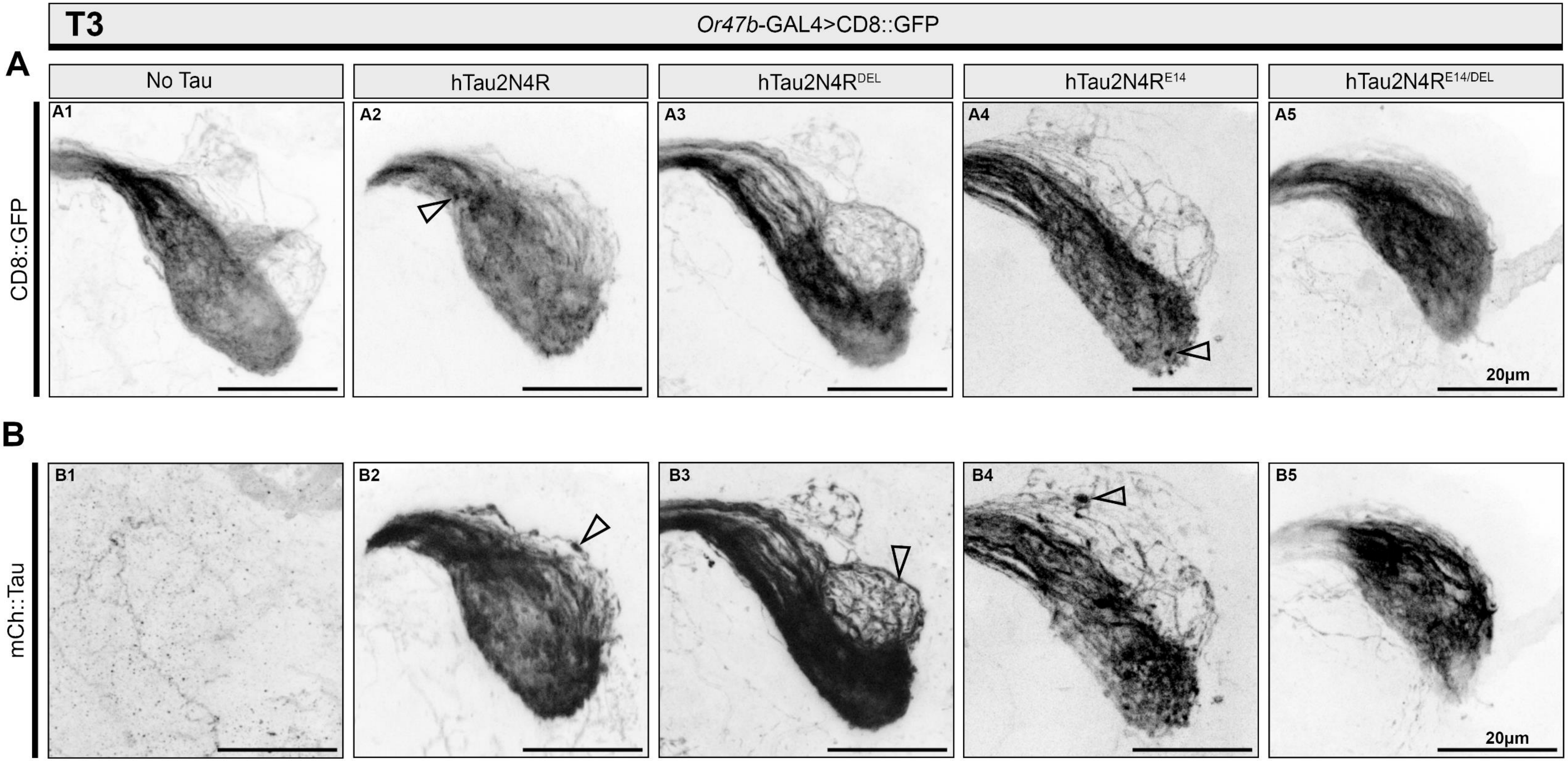

**Supplementary Figure 5**

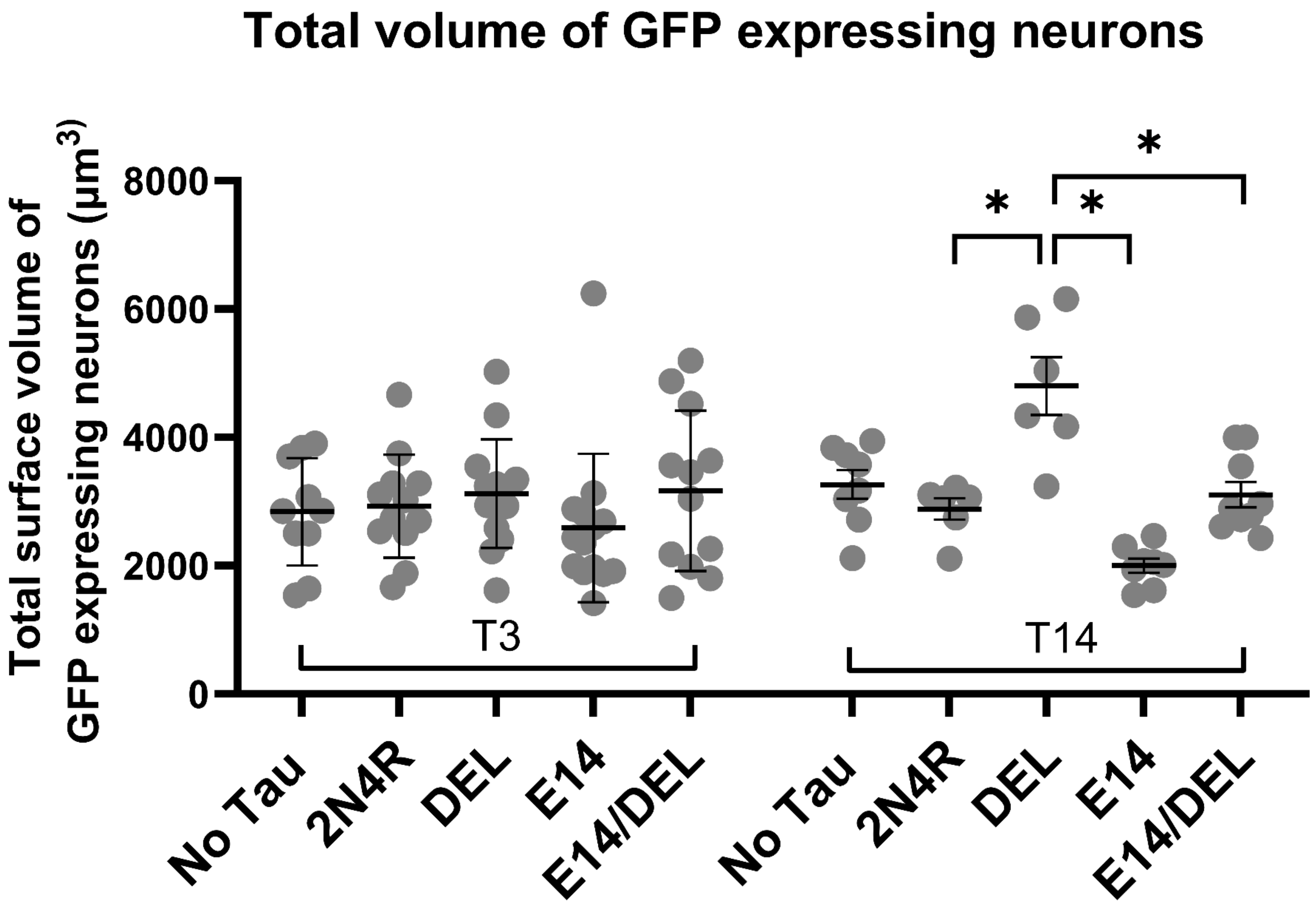

**Supplementary Figure 6.**

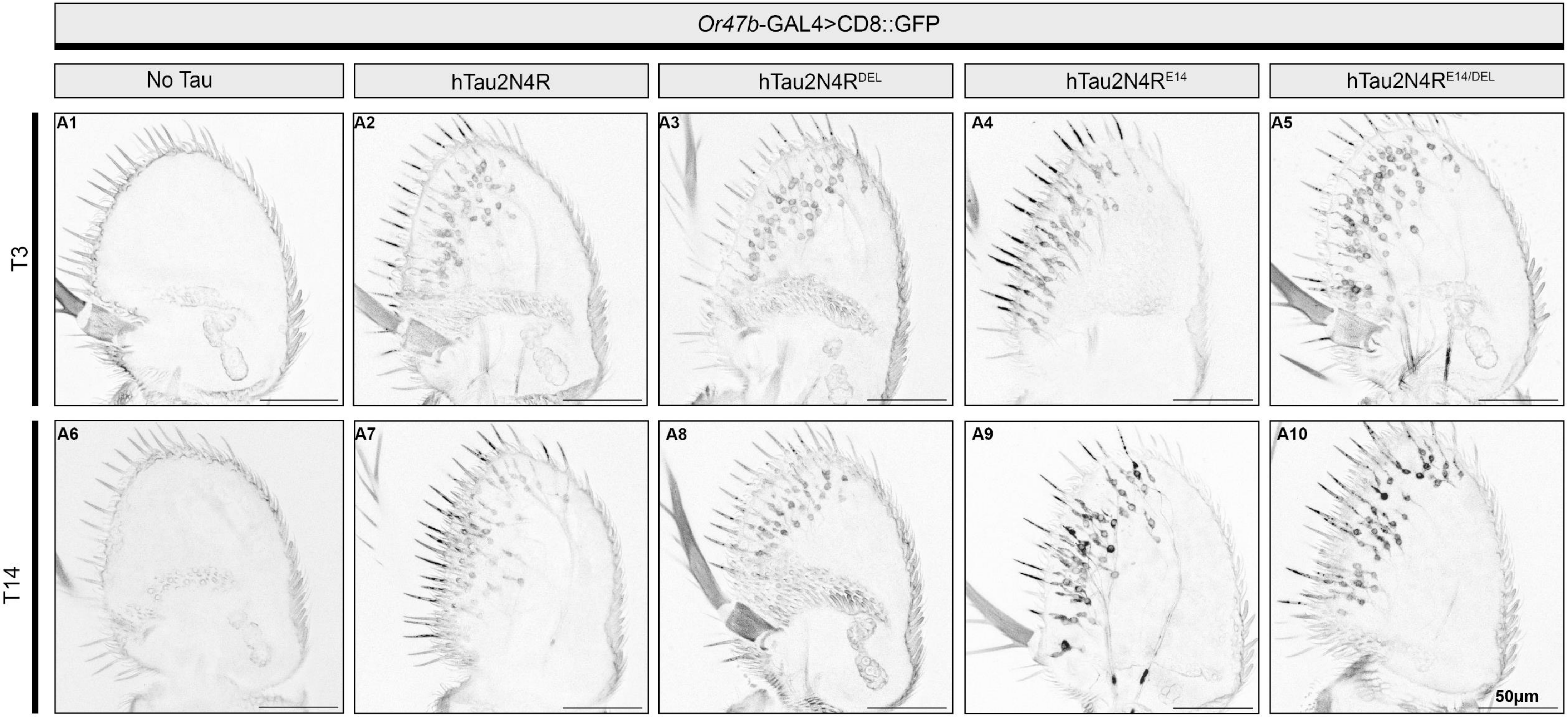

**Supplementary Figure 7.**

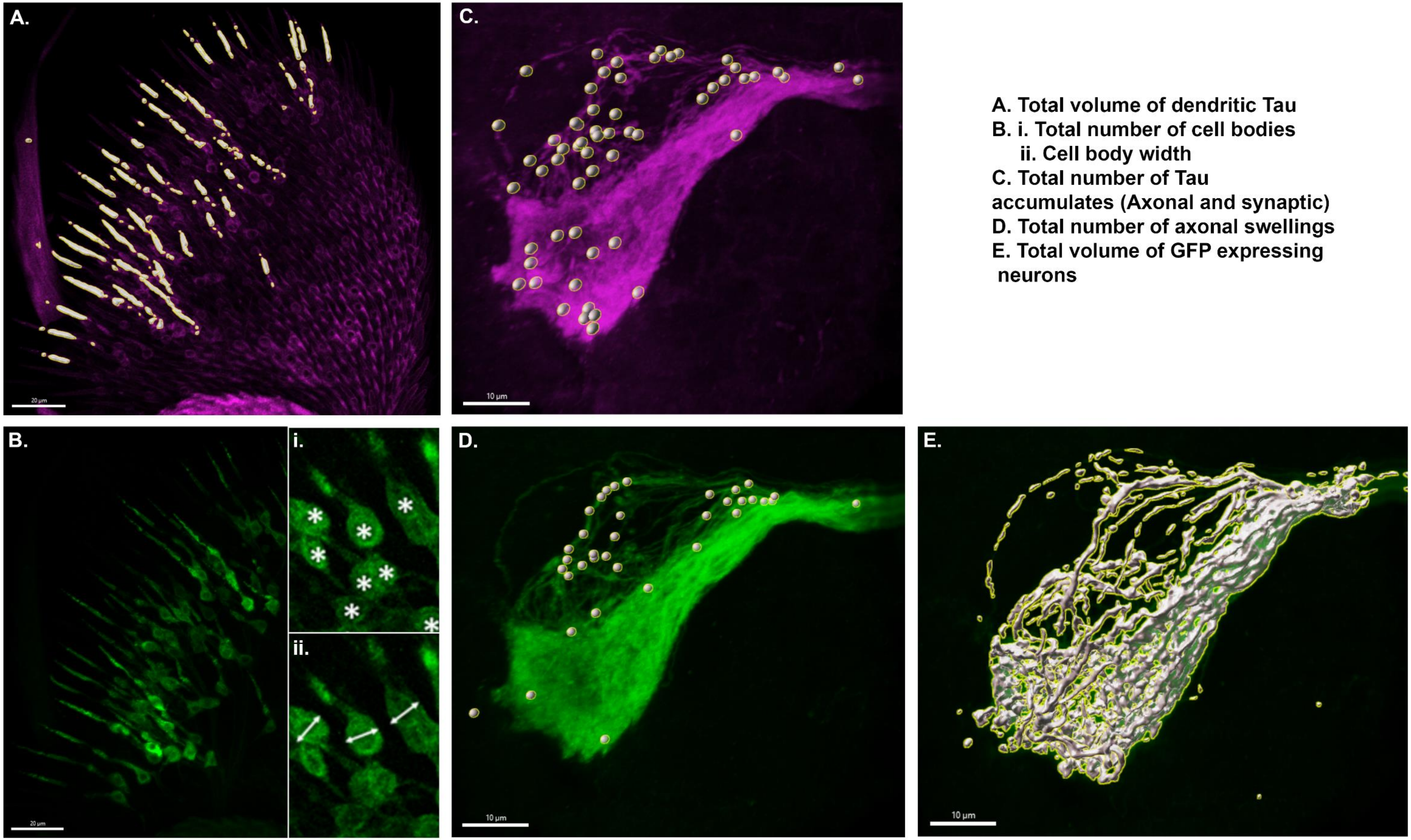


A. Total volume of dendritic Tau
B. i. Total number of cell bodies
    ii. Cell body width
C. Total number of Tau accumulates (Axonal and synaptic)
D. Total number of axonal swellings
E. Total volume of GFP expressing neurons
